# *Hox* genes regulate asexual reproductive behavior and tissue segmentation in adult animals

**DOI:** 10.1101/2021.01.24.427972

**Authors:** Christopher P. Arnold, Analí Migueles Lozano, Frederick G. Mann, Jeffrey J. Lange, Christopher Seidel, Alejandro Sánchez Alvarado

## Abstract

*Hox* genes are highly conserved transcription factors renowned for their roles in the segmental patterning of the embryonic anterior-posterior (A/P) axis^1^. Emerging evidence for *Hox* gene expression and function in postnatally derived structures has fueled interest in their additional roles beyond embryogenesis^2,3^. We report novel functions for *Hox* genes in A/P adult tissue segmentation and transverse fission behavior underlying asexual reproduction in the planarian flatworm, *Schmidtea mediterranea.* Silencing of each of the planarian *Hox* family members identified 5 *Hox* genes required for asexual reproduction. Among these, silencing of *hox3* genes resulted in supernumerary segments, while silencing of *post2b* eliminated segmentation altogether. The opposing roles of *hox3* and *post2b* in segmentation are paralleled in their respective regulation of fission behavior. Silencing of *hox3* increased the frequency of fission behavior initiation, while silencing of *post2b* eliminated fission behavior entirely. Furthermore, we identified a network of downstream effector genes mediating *Hox* gene regulation of asexual reproduction, thereby providing insight into their respective mechanisms of action. Our study establishes postembryonic roles for *Hox* genes in regulating the emergence of tissue segmentation and specific behaviors associated with asexual reproduction in adult animals.

---

We recently uncovered and characterized the size-dependent development of adult tissue structures underlying asexual reproduction in the planarian, *Schmidtea mediterranea*^*4*^ (**Fig1a**). These highly regenerative flatworms undergo transverse fission – an asexual reproductive behavior in which torn posterior tissue fragments regenerate and give rise to clonal progeny^5^. While historically designated as unsegmented flatworms, our study revealed a cryptic form of segmentation that antero-posteriorly allocates tissue for fission progeny in coordination with adult animal growth^*4*^. In concert with this segmentation process, differential patterning of the planarian central nervous system (CNS) establishes and modulates fission behavior in accordance with animal size^*4*^. The molecular mechanisms mediating and coupling these parallel size-dependent processes are still unclear. *Hox* genes encode transcription factors with conserved roles in the regulation of A/P developmental patterning^1^. Emerging evidence indicates additional roles for *Hox* genes in the segmentation of developing tissues as well functions in the formation of postnatal tissue structures^2,6^. We therefore hypothesized that *Hox* genes play a role in A/P directed tissue segmentation and/or transverse fission behavior underlying asexual reproduction in adult animals.

**Fig. 1:**
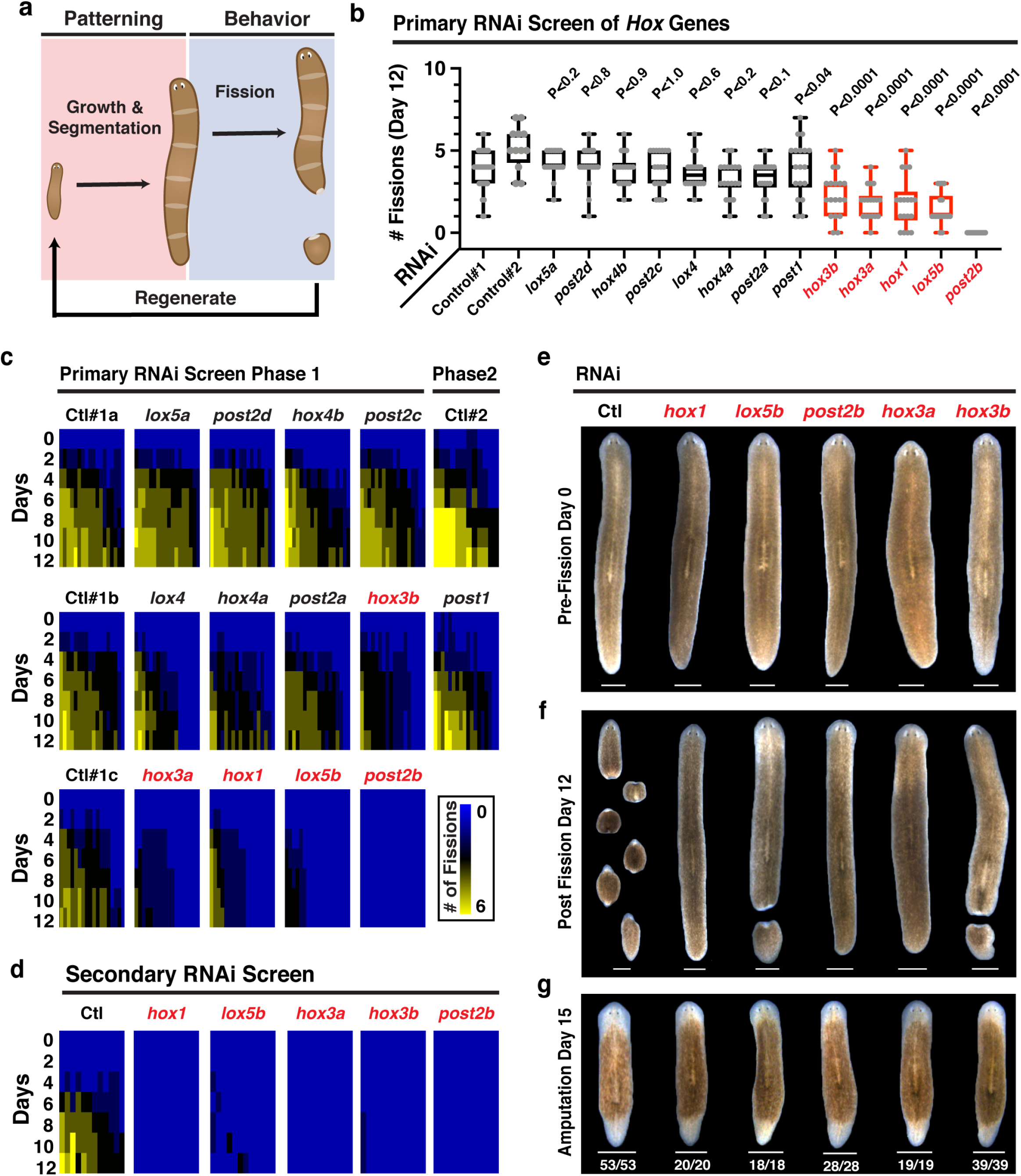
Planarian *Hox* genes are required for adult asexual reproduction. **a,** Diagram of the patterning and behavioral aspects of planarian asexual reproduction. **b,** Fission activity following RNAi against *Hox* genes. Plot depicts fission progeny number on day 12 of primary RNAi screen (n=12, 18, or 54 animals; p-value calculated by Welch’s two-tailed t-test versus corresponding control). **c, d,** Heatmaps depicting fission activity following RNAi treatment for both the (**c**) 2-phase primary and (**d**) secondary *Hox* RNAi screens. Cumulative fissions over time are displayed for individual worms from each RNAi condition (n=12 or 18 animals). **e, f** Representative images of animals from secondary *Hox* RNAi screen on days 0 and 12 of the fission assay (n= 12 animals, 3 independent repeats). Animals were given 9 dsRNA feedings for primary phase I and 12 dsRNA feedings for phase II screening. Animals were given 17 bacterial RNAi feedings for secondary screenings. **g,** Representative images and phenotypic frequency of RNAi animals 15 days post amputation after 8 dsRNA feedings (n=18-53 animals, 3 independent repeats). Scale,1mm.

The planarian genome encodes 13 *Hox* genes with conserved homeodomains: *Hox1*, *Hox3a*, *Hox3b*, *Hox4a*, *Hox4b*, *Lox4*, *Lox5a*, *Lox5b*, *Post1*, *Post2a*, *Post2b*, *Post2c*, *Post2d^7^* (**ED Fig1a, b**). Planarian *Hox* genes are dispersed throughout the genome in an atomization of the ancestral cluster^1^ (**ED Fig1c)**. Their expression generally increases from embryogenesis to adulthood (10/13 genes) with comparatively minor changes during whole animal regeneration/remodeling^8,9^ (**ED Fig2, 3**). Whole mount *in situ* hybridization (WISH) of adult stage planarians indicates that a subset of *Hox* gene transcripts are axially restricted (*hox3b*, *hox4b*, *lox5a*, *post2c*, and *post2d*) while others are radially expressed (*post2a* and *post2b*)^7^. Despite an abundance of transcriptional analysis and considerable experimental effort, the functions of *Hox* genes in planaria have remained largely a mystery^7,10,11^.

We determined the role of each of the planarian *Hox* genes in asexual reproduction. Individual RNAi knockdown of 5/13 *Hox* genes (*hox1*, *hox3a*, *hox3b*, *lox5b*, and *post2b*) in adult stage planaria resulted in a greater than 50% reduction in the generation of fission progeny (**Fig 1b,c**, **ED Fig4a-c**). Additional rounds of *Hox* gene knockdown in a secondary screening either completely or nearly completely eliminated fission activity (**Fig 1d, f, ED Fig4d**). Consistent with previous reports, knockdown of *hox1*, *lox5b*, *post2b*, *hox3a*, or *hox3b* did not result in evident morphological changes during growth, homeostasis, or regeneration^7^ (**Fig 1e-g**). These results identify a set of planarian *Hox* genes with prominent roles in asexual reproduction.

We hypothesized that planarian *Hox* genes regulate the adult tissue segmentation underlying asexual reproduction. We utilized our previously established physical compression assay to reveal the fission planes dividing tissue segments in RNAi animals, and thereby determine the respective effects of *Hox* gene silencing^4^. Knockdown of *hox1* or *lox5b* had little to no effect on segment number (**Fig2a-b**). In contrast, knockdown of *post2b* eliminated fission segmentation entirely, while knockdown of *hox3a* or *hox3b* resulted in supernumerary segments (**Fig2a-b**). Although simultaneous knockdown of *hox3a*+*hox3b* increased fission segment number, it also reduced fission progeny yield (**Fig2a-b, ED Fig4e-g**). Increasing the number of fission planes dividing the animal reduces the size of each segment and the resultant fission progeny. We quantified the size of rare fission progeny in *hox3* RNAi animals, revealing a decrease in progeny size consistent with the increase in segmentation observed via physical compression (**Fig 2c-d**). Altogether, these data indicate that the planarian *Hox* genes *post2b* and *hox3* mediate the number of adult A/P tissue segments associated with asexual reproduction.

**Fig. 2:**
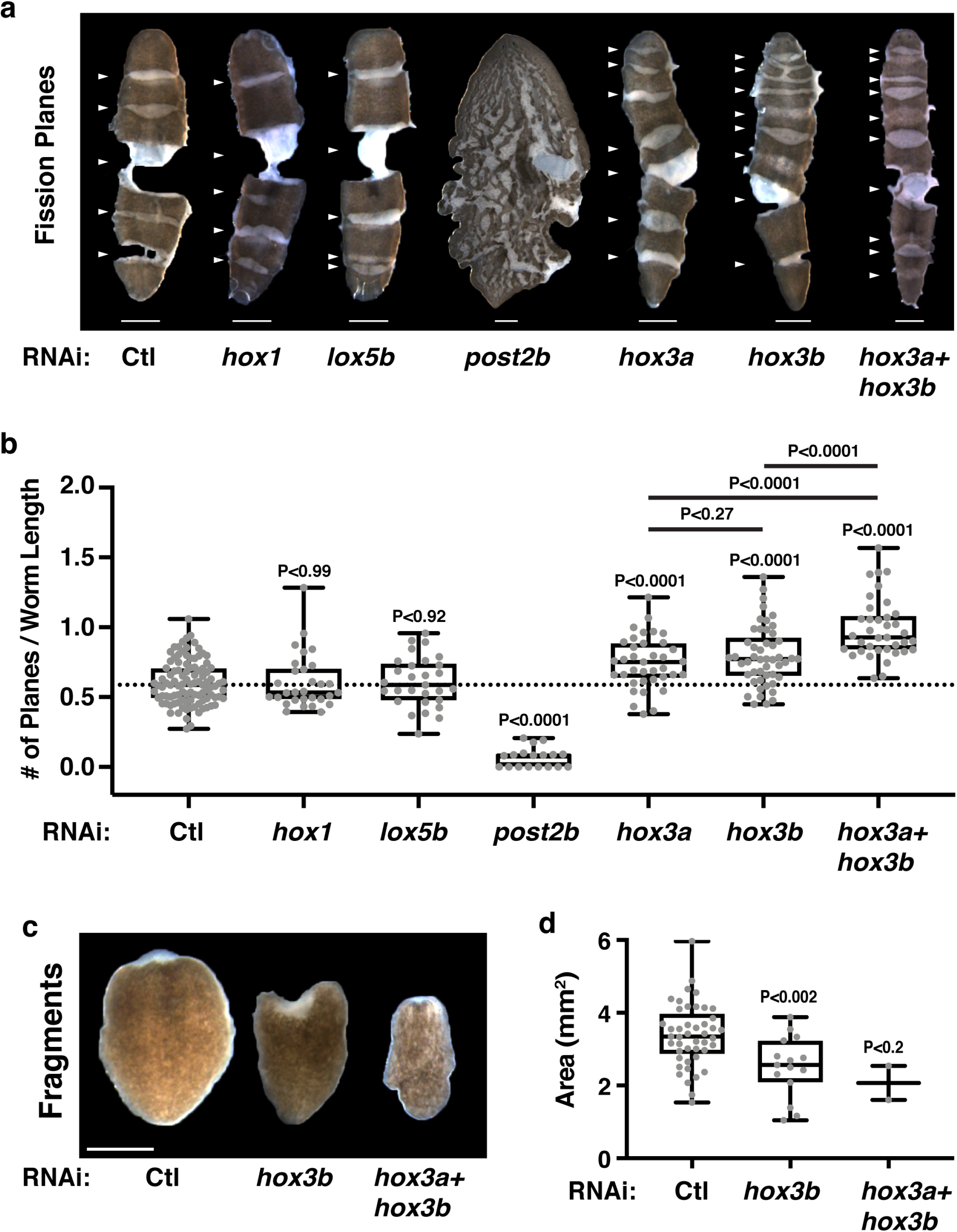
*Post2b* and *Hox3* are opposing regulators of adult tissue segmentation. **a,** Representative images of RNAi animals compressed to reveal fission planes and segmentation (arrows indicate fission planes). **b,** Plot of the number of fission planes/animal length following *Hox* RNAi treatment (n=18-101 animals pooled from two independent experiments with 8 dsRNA feedings or 17 bacterial RNAi feedings). **c,** Representative images of fission progeny following RNAi treatment. **d,** Plot of surface area of fission progeny (n=2-45 fission progeny, 17 bacterial RNAi feedings). p-values calculated by Welch’s two-tailed t-test. Scale, 1mm.

We next determined the roles of planarian *Hox* genes in the frequency, duration, and success of asexual reproductive behavior. Using automated webcam time-lapse imaging, we recorded fission events in *Hox* RNAi treated animals^4^ (**Fig 3a, ED Fig5, 6, Videos S1-8**). Individual knockdown of either *hox1* or *lox5b* substantially reduced the success of fission behavior (**ED Fig 6b-f, I, Video S2, S5**). Mirroring their opposing roles in segmentation, *post2b* RNAi eliminated fission behavior entirely (**ED Fig 6j, 7b**) while *hox3a*+*hox3b* RNAi increased fission behavior frequency and duration (**Fig 3b-d, ED Fig 6g-h, 7a, c, Video S6, S7**). Individual RNAi of *hox3b,* but not *hox3a,* largely phenocopied *hox3a*+*hox3b* RNAi effects on fission behavior (**ED Fig6b-d, g, h, ED Fig 7c, Video S3, S4**). These results suggest unequal levels of functional redundancy for the *hox3* genes in the regulation of fission behavior in contrast to their similar contributions to fission segmentation (**Fig 2**). Importantly, these results reveal that *post2b* and *hox3* mediate opposing roles in the regulation of fission behavior that parallel their roles in the segmentation of adult tissues.

**Fig. 3:**
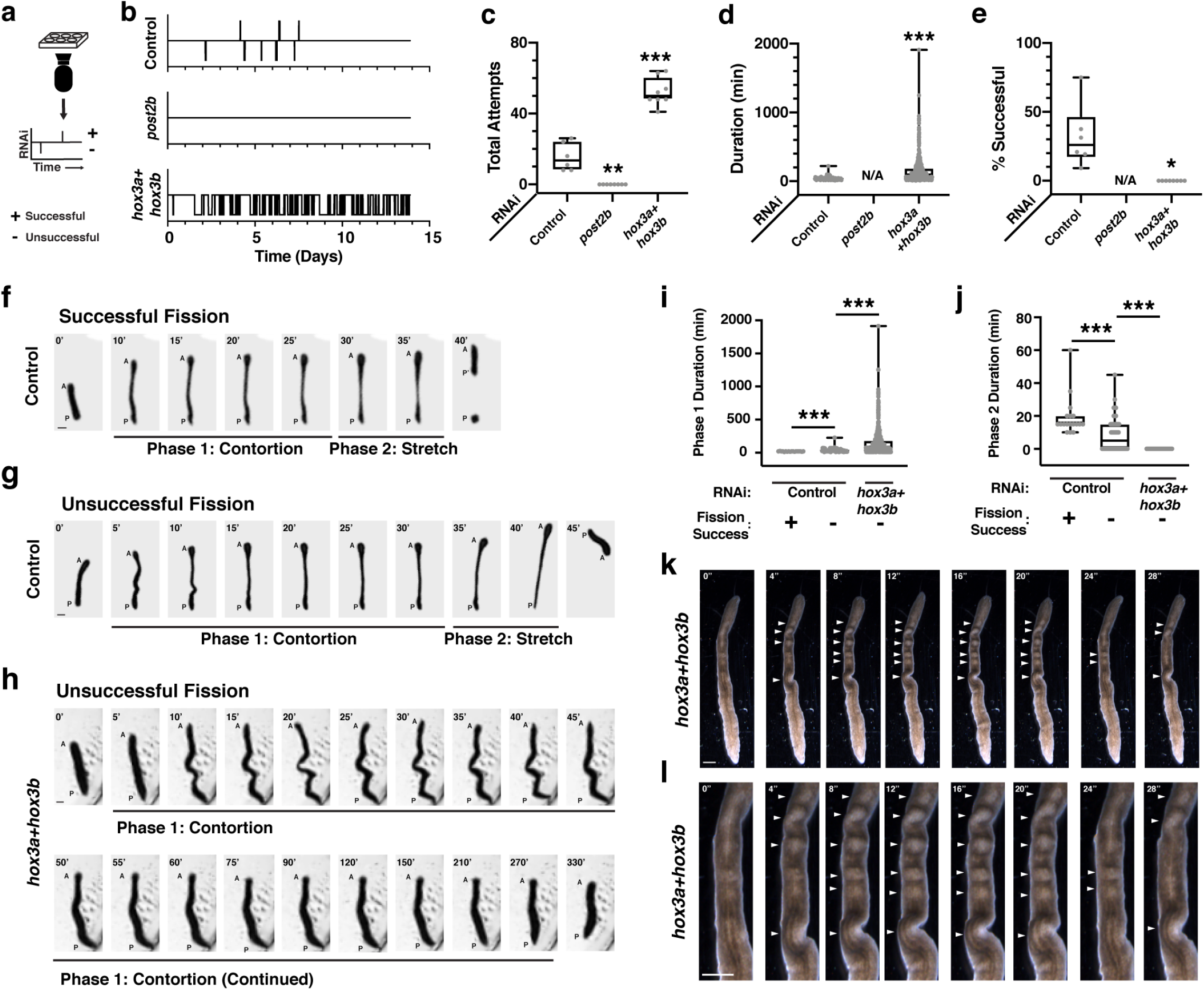
*Post2b* and *Hox3* are opposing regulators of asexual reproductive behavior. **a,** Diagram of webcam time-lapse imaging and timeline visualization depicting successful (upward displacement) and unsuccessful (downward displacement) fission attempts. **b,** Representative activity timelines of fission behavior from live imaging of control, *post2b*, and *hox3a*+*hox3b* RNAi animals. **c-e,** Plots of (**c**) total number of fission attempts, (**d**) percentage of successful attempts, and (**e)**duration of each attempt for RNAi treated animals (n=6-8 animals) **f-h,** Representative images (5 minutes per frame) of (**f**) successful and (**g, h**) unsuccessful fission attempts divided into Phase 1 (Contortion) and Phase 2 (Stretch) for (**e, f**) control and (**g**) *hox3a*+*hox3b* RNAi-treated animals. **i, j,** Plots of the duration of (**i**) Phase 1 and (**j**) Phase 2 within each fission attempt for RNAi treated animals (n=6-8 animals). **k**, **l**, Representative images of the (**k**) entire body or the (**l**) pre-pharyngeal region of *hox3a*+*hox3b* RNAi-treated animal stalled in Phase 1 (n=10/10 animals, arrows indicate tissues undergoing peristaltic contractions). Animals given 17 bacterial RNAi feedings. Scale, 1mm. p-values calculated by Welch’s two-tailed t-test.

Our analysis indicates that planarian *hox3* genes have shared roles in the negative regulation of adult tissue segmentation and asexual reproductive behavior. Yet, while *hox3* RNAi increases the frequency and duration of fission behavior, these attempts are ultimately unsuccessful (**Fig3b-e**). We hypothesized that *hox3* is also required at a later stage of the asexual reproduction process. Detailed analyses of fission behavior in control and *hox3* RNAi-treated animals revealed two distinct phases of behavior (I & II) common to individual fission attempts (**Fig3f, g**). During phase I, animals elongate anteriorly in a contorted manner. During phase II, animals further stretch out, thinning and rupturing the fission plane tissue and releasing the progeny. Analysis of *hox3* RNAi animals reveals an inability to progress from phases I to II, thus preventing completion of asexual reproduction (**Fig3h-j**). Furthermore, high resolution time-lapse imaging of *hox3a*+*hox3b* as well as *hox3b* RNAi revealed that phase I contortions are coincident with A/P directed peristaltic contractions (**Fig3k, l, Video S9, S10**). This peristalsis was previously observed in live imaging of wildtype animals, but its significance in planarian asexual reproduction is still unclear^4^. We conclude from these data that *hox3* is required for both the initiation and successful progression of asexual reproductive behavior.

Our findings demonstrate that *Hox* genes regulate asexual reproduction via the emergence and modulation of A/P segmentation and behavior in adult animals. *Hox* genes mediate their functions via direct and indirect regulation of downstream effector genes^12^. To elucidate planarian *Hox* downstream effector genes regulating adult tissue functions, we devised a 2-part strategy to (I) identify putative candidates via differential gene expression analysis, and (II) functionally validate their roles in fission (**Fig4a**). RNAseq analysis identified 1,724 differentially expressed genes (DEGs) following perturbation of *Hox* gene function (**Fig4b**). We selected and successfully cloned 423 putative effector genes into RNAi vectors for functional analysis. We determined the role of each gene in asexual reproduction by scoring resultant fission progeny following RNAi-mediated gene perturbation (**ED Fig8a, b**). We focused on RNAi conditions that phenocopied *Hox* gene RNAi - i.e., robustly decreased fission progeny (>4-fold reduction, p-value<0.01) independent of evident alterations to body plan morphology or size (**Fig1e-g, ED Fig4d**). Using these criteria, our functional screen identified 24 putative *Hox* downstream effector genes required for asexual reproduction (**Fig4c, d, ED Fig8c, d, Table 1).**

**Fig. 4:**
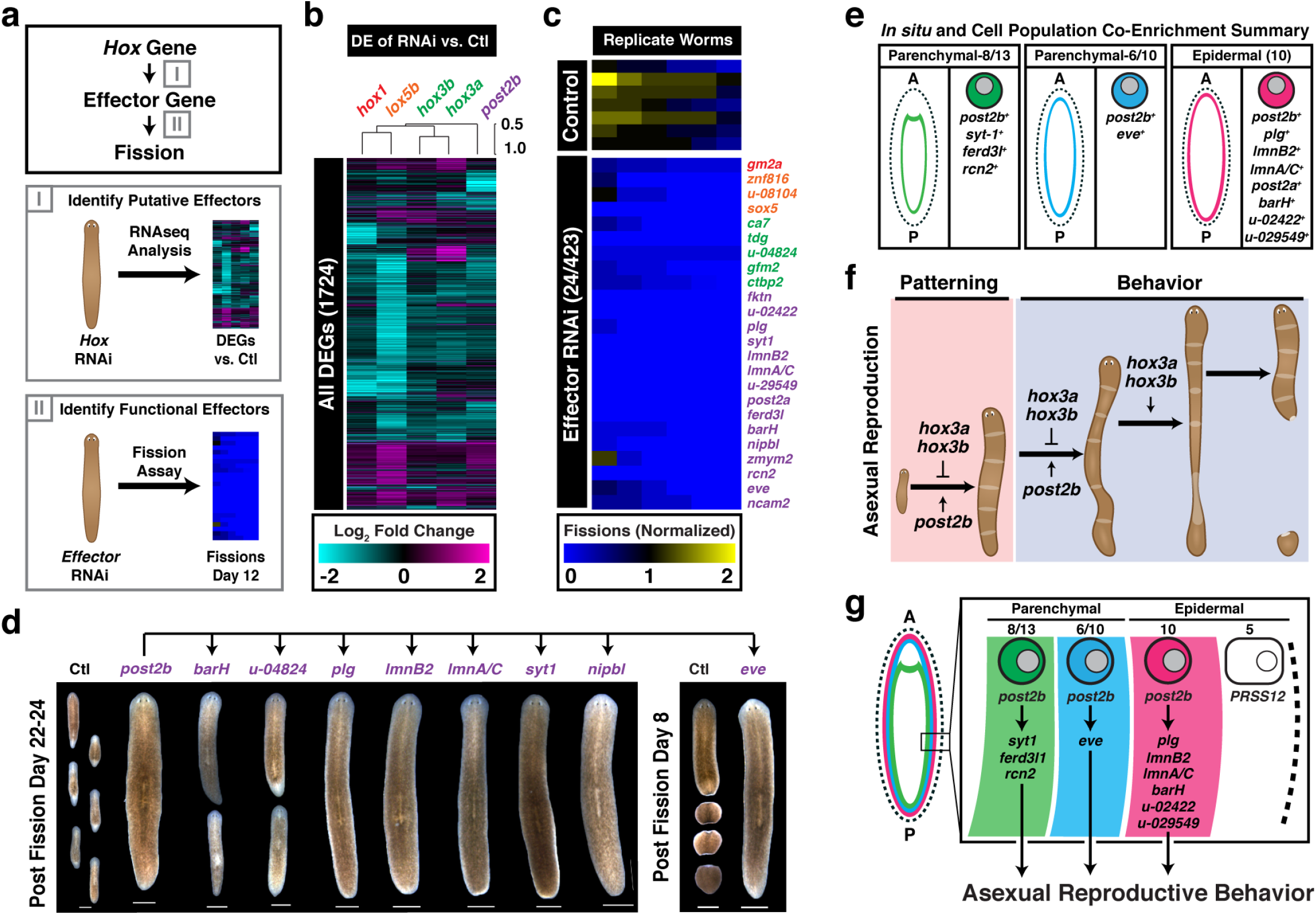
Identification of downstream effectors mediating *Hox* gene regulation of asexual reproduction. **a,** Diagram depicting a 2-part strategy to elucidate downstream effectors of *Hox* genes that regulate fission: (I) use RNAseq analysis to identify putative effector genes from the differentially expressed genes in *Hox* RNAi-treated animals, then, (II) use RNAi to determine the effects of putative effector genes on fission. **b,** Heatmap of 1724 DEGs from RNAseq that are significantly regulated following RNAi of *Hox* genes. **c,** Heatmap of the top 24/423 genes from RNAi screen of putative downstream fission effectors (See supplemental figure 8, 16–19 Bacterial RNAi feedings). Heatmap depicts normalized fission number on Day 12 across six RNAi-treated worms. **d,** Representative images of *post2b* effector RNAi-treated animals 22–24 days or 8 days post fission induction. Scale, 1mm. **e,** A/P location (left subpanel) and cell population specific enrichment (right subpanel) for the transcripts of *post2b* and its downstream effectors (summary of data from the cell type transcriptome atlas Fincher et al, digiworm.mit.edu). Depicted are populations from the radial epidermal and parenchymal lineages (10/11 *post2b* effectors). **f,** Diagram summarizing the roles of the *Hox* genes *post2b*, *hox3a*, and *hox3b* as regulators of the adult tissue patterning and behavior required for asexual reproduction. **g**, Model depicting the mechanism by which *post2b* regulates asexual reproductive behavior via the direct and/or indirect regulation of downstream effectors within the radial cell populations.

The uncovered effector genes include putative mediators of *hox1*, *lox5b*, *hox3*, *hox3b*, *hox3a*, and *post2b* gene function in asexual reproduction (**Fig8c, Table 1**). Of note, we identified *post2a* as a downstream effector gene of *post2b*. Effects of *post2a* RNAi in our primary screen were near significance (p<0.1), but extension of RNAi knockdown (8X vs. 17X RNAi feedings) increases phenotypic penetrance, yielding significant effects on fission (**Fig1b, Fig4c, d**). Of the remaining effector genes, 20/23 have homology to human genes, indicating that planarian *Hox* genes regulate asexual reproduction by controlling conserved homologous genes rather planarian-specific genes (**Table 1**). Furthermore, the identification of a *Hox* downstream effector gene homologous to *ncam2* is significant as N-CAM was the first identified mammalian *Hox* gene target^13^. Analysis of the 1,000bp upstream regulatory region reveals the presence of conserved *Hox* binding motifs (TTWATKA) in 22/24 effector genes, including *ncam2* (**Table 1**). These findings suggest that planarian asexual reproduction may be mediated by conserved *Hox* gene/target gene regulatory interactions and illustrate the potential of this novel paradigm for resolving conserved mechanisms of action underlying *Hox* gene function.

In order to gain insight into the mechanism of action by which *Hox* genes regulate asexual reproduction, we set out to determine the spatial and cellular context of *Hox* genes and their effectors. *In situ* hybridization analysis of *Hox* gene transcripts poses a significant challenge due to their low expression in planarian tissues^7^. We therefore utilized published scRNAseq expression data to resolve the cell populations in which *Hox* genes were co-expressed with their cognate downstream effectors^14^. Using this approach, we resolved overlapping expression for *post2b* and 10/15 of its downstream effectors (**Table 1**). *Post2b* is enriched in the *smedwi*^+^-PP, brain-PP1, and the non-ciliated neurons-0 cell populations in addition to the radial and medio-laterally layered parenchymal-8/13, parenchymal-6/10, and epidermal-10 cell populations (**Fig4e, ED Fig8e**). Amongst the 10 resolved effector genes, *rcn2* was co-enriched in the *smedwi*^+^-PP, brain-PP1, and parenchymal-8/13 populations (**Fig4e, ED Fig8e, Table 1**). The remaining 9 effectors were co-enriched in the radially layered parenchymal/epidermal cell populations: *syt-1* and *ferd3l* in parenchymal-8/13, *eve* in parenchymal-6/10, and *plg*, *lmnB2*, *lmnA/C*, *post2a*, *barH*, *unk-02422*, and *unk-029549* in epidermal-10 (**Fig4e, Table 1**). To gain insight into the function of these *post2b* effectors, we determined whether they were required for segmentation similar to *post2b*. Surprisingly, we found no evidence to support that these radially expressed *post2b* effectors function in segmentation (**ED Fig8f**, data not shown). We postulate that effector function in these radial cell populations is instead related to functions within the nervous system underlying asexual reproductive behavior.

*Hox* genes have previously established roles in synaptic specificity, neuronal subtype specification, and selective muscle innervation underlying vital behaviors^15^. Additionally, radially expressed *post2b* effectors are homologous to genes with functions in the nervous system. The effectors *rcn2* and *syt1* encode genes homologous to binders and sensors of intracellular calcium release, a key regulator of rhythmic muscle contraction and synaptic transmission^16,17^. *Post2b* effectors also include genes homologous to a bHLH transcription factor (*ferd3l1)* and two homeobox transcription factors (*barH* and *eve*) that regulate neuronal development ^18–20^. Additionally, nuclear lamins (homologous to *lmnA/C* and *lmnB2*) are required for neural circuit integrity and neuronal migration^21,22^. In summary, *post2b*/effector genes are required for fission behavior, exhibit cell population specific expression, and have homology to genes with previously reported roles in neuronal function. These results argue that the identified radial cell populations constitute a cellular basis for *Hox* mediated regulation of asexual reproductive behavior with likely functions in the nervous system. *Hox* genes have well established embryonic roles in the specification of segmental identity across the A/P axis of the body plan and within sub-structures of the nervous system such as the vertebrate hindbrain, a complex coordinator of motor activity and rhythmic behaviors^1,23^. Our work demonstrates a novel adult-stage function for *Hox* genes in the coupling of body-wide tissue segmentation with specific animal behaviors.

In conclusion, our study identifies roles for planarian *Hox* genes as mediators of adult tissue segmentation and behavior, as well as providing the first evidence that *Hox* genes function in asexual reproduction (**Fig4f, ED Fig 8g, h).** Recently, the discovery that the expression of homeodomain proteins constitutes a unique code delineating all of the neuron classes of *C. elegans* has provided further evidence for the prominent roles of *Hox* family members and other homeobox genes in the evolution of neuronal cell type specification^24^. Our study expands upon this body of work by identifying functional roles for *Hox* genes in the establishment and regulation of specific behaviors in adult animals. Our current findings in combination with prior work indicate complementary roles for *Hox* and *Wnt* genes during planarian asexual reproduction^4^. As adult animals grow, the collective activities of *hox1*, *hox3a*, *hox3b*, *lox5b*, *post2*a and *post2b* are required for the establishment and regulation of asexual reproductive tissue segmentation and behaviors. In parallel, Wnt activity maintains A/P identity of the body plan^25^, and coordinates growth with size-dependent patterning of fission modulatory mechanosensory neurons^4^. Finally, re-establishment of a gradient of Wnt activity modulates gene expression (including activation of the axial *Hox* genes *hox4b*, *lox5a*, *post2c*, and *post2d*) to regenerate the fissioned posterior tissue fragment into a clonal progeny^10^. Together, these findings suggest that a subset of Wnt-independent *Hox* genes mediate specific functions in asexual reproduction and work in concert with Wnt-dependent patterning and regeneration programs. From cnidarians to deuterostomes, *Hox* genes play conserved roles in the regulation of axial patterning^1,6^. Notably asexual reproduction, via budding or fission, is also distributed throughout these groups and phyla^26^. We speculate that the regulation of asexual reproduction and/or the specification of neural circuitry utilized in this process represents an ancestral function for *Hox* genes.

## Acknowledgements

We thank members of the ASA laboratory for discussion and advice, and Robb Krumlauf for comments. We are grateful to the Stowers Planarian and Molecular Biology core facilities for technical contributions and methods development.

## Data availability

Original data underlying this manuscript can be accessed from the Stowers Original Data Repository at http://www.stowers.org/research/publications/LIBPB-1567

## Authors contributions

Conceptualization, data analysis, and interpretation (CPA, ASA), acquisition of data (CPA, AML, JJL), design, fabrication, and software for planarian live imaging systems (JJL), cloning of RNAi vector library for gene perturbation and screening (FGM), RNAseq Differential Gene Expression Analysis (CS), writing of the original manuscript (CPA), supervision and funding acquisition (ASA), and revision and editing of the manuscript (all authors).

## Materials and Correspondence

Correspondence should be directed to

## Methods

### Animal Husbandry

Clonal CIW4 *Schmidtea mediterranea* were maintained in 1X Montjuic salts as previously described^4^. CIW4 animals were sourced from a large recirculation culture and placed placed directly into to a unidirectional flow system culture for RNAi feeding experiments as previously reported^4,27^

### Gene cloning and RNAi Feeding protocol

Gene cloning was performed as previously describe but with the following distinctions^4^. Primer3 was used to design primer pairs targeting genes of interest, using the Sánchez Alvarado lab transcriptome as a reference. 5’ overhangs were added to ends of both forward and reverse primers to enable downstream Gibson Assembly. Amplified products were designed to be 400–600 base pairs in length. PCR was performed using cDNA as template. Products were inserted into a modified version of the pPR-T4P vector and transformed directly into *Escherichia coli* strain HT115. Final cloning products were verified by sequencing. Preparation of dsRNA and RNAi feedings were performed as previously described with the following distinctions^4^. Purified dsRNA food was prepared by mixing 1 volume of dsRNA at 400ng/ml with 1 volume of beef liver paste. Bacterial RNAi food was prepared by mixing 8ml of pelleted bacteria with 120ul of beef liver. Animals were fed from a starting size of less than 3mm to a final size of greater than 10mm. To achieve this, worms were either fed 7–9 times with purified dsRNA food or 16–19 times with bacterial RNAi food as indicated. Control animals were fed RNAi food targeting *unc22*.

### Fission Assay

Fission induction was performed as previously described with the following distinctions^4^. Individual animals into the wells of a 6 well tissue culture plate. The number of fission fragments was scored every 1 to 2 days. For data analysis, the number of daily cumulative fissions was directly plotted normalized to the average of the control RNAi fissions where indicated. Fission number was converted to a heat color code for visualization.

### Fission Segmentation Compression Assay

Fission segmentation was revealed by compression of worms between a glass coverslip and plastic tissue culture dish as previously described^4^.

### Microscopy

Images of live worms and regenerating fission/amputation fragments were acquired using a Leica M205 microscope. Fission fragment area was calculated using Adobe Photoshop.

### Live Imaging of Fission Behavior

High-throughput imaging of fission behavior was performed as previously described, but with the following modifications. For all data sets, worms were placed in 6 well dishes and imaged in one of two ways. For time-lapse A (**ED Fig, S6**, **Video S1-5)**, worms were imaged from above with a single 4k webcam (Logitech Brio). Four inverted LED ring lights (AmScope) mounted above the camera were used for illumination. Images were acquired using an in house written Python script described previously^4^. In time-lapse B (**Fig3**, **ED Fig7**, **Video S6-8**), images were acquired with a new configuration where the camera was mounted below a large (2ft x 2ft) plexiglass panel that was topped with large white plexiglass panels. The same 4 LED lights were placed at the bottom of the set-up next to the camera. In this format a webcam (Logitech c920) was used to acquire images. The time-lapse was acquired using micro-manager (https://micro-manager.org/) and the opencvgrabber framework. In both cases, the camera gain, exposure, and auto focus were controlled using the Logitech Webcam Controller software (https://download01.logi.com/web/ftp/pub/video/lws/lws280.exehttps://download01.logi.com/web/ftp/pub/video/lws/lws280.exe).

Videos of individual animals were analyzed, manually annotated, and example frames were extracted using Image J^4^. Fission attempt initiation was scored at the frame in which a rapid stationary elongation occurred following a period of contraction and immobility. Termination of fission attempt was scored at the frame in which the animal ruptured and released a fragment (successful) or the animal returned to its original pre-elongation state (unsuccessful). Fission events were divided into phase I (initial elongation accompanied by body contortions) and phase II (second elongation accompanied by thinning and/or rupture). Data was visualized with timelines as previously described^4^. Worms that crawled out the well and/or desiccated during the course of the experiment were not considered for further analysis.

Peristaltic activity of *hox3b* and *hox3a*+*hox3b* RNAi worms was captured using a Leica M205 microscope (acquired at 10 frames per second, **Video S9,10**).

### RNAseq Analysis

Ctl or *Hox* gene RNAi animals (7x purified dsRNA feedings) animals were starved 5 days and snap frozen for RNA collection (4 samples with 3 animals each). RNA was extracted using Trizol Reagent and Dnase treated (Qiagen Kit Cat#) as previously described^27^. Libraries were prepared according to manufacturer’s instructions using the TruSeq Stranded mRNA Prep Kit (Illumina). The resulting libraries were purified using the Agencourt AMPure XP system (Beckman Coulter) then quantified using a Bioanalyzer (Agilent Technologies) and a Qubit fluorometer (Life Technologies). Libraries were re-quantified, normalized, pooled and sequenced on an Illumina NextSeq 500 instrument as High Output 75bp single read runs using NextSeq Control Software version 2.2.0.4. Following sequencing, Illumina NextSeq Real Time Analysis version 2.4.11 and bcl2fastq2 version 2.20 were run to demultiplex reads and generate FASTQ files. These FASTQ files were then aligned to the smed_dd_g4 genome using STAR and parameters: --outFilterMultimapNmax 2 --alignIntronMax 5000 --quantMode GeneCounts, along with the Smes_hv_v1 gene models from the Max Plank Institute, obtained via Planmine, to generate read counts to genes^28^. Differentially expressed genes were determined using the edgeR library in R^29^. P-values were adjusted for multiple hypothesis testing by the method of Benjamini and Hochberg^30^. The data have been deposited in GEO with accession number: GSE159876.

### Planarian Hox Gene, Transcript, and Protein Sequence Analysis

Planarian Hox gene mRNA and amino acid sequences were obtained from Planosphere (https://planosphere.stowers.org/find/genes).). The genomic contig location of planarian Hox genes was determined using PlanMine v3.0 (http://planmine.mpi-cbg.de/planmine/blast.do).). Homeodomains of planarian *Hox* genes were annotated using Pfam 33.1 (https://pfam.xfam.org/)^31^.. Homeodomain sequences were aligned via Clustal Omega (https://www.ebi.ac.uk/Tools/msa/clustalo/).

### Planarian Hox Gene and Effector Gene Expression

Expression of Hox genes during regeneration and development were obtained from Zeng et al., 2018 and Davies et al., 2017, respectively^8,9^. Cell population enrichment of transcripts and representative cell population in situ patterns obtained from a planarian cell transcriptome atlas (https://digiworm.wi.mit.edu/)^14^..

### Data Visualization and Statistical Tests

All heatmaps were generated using Multiple Experiment Viewer (MeV). All graphs were generated using GraphPad Prism. All illustrations were made using Adobe Illustrator. For pair-wise comparisons, significance was calculated with an unpaired t test with Welch’s correction using GraphPad Prism.

## Extended Data Figure Legends

**Extended Data Figure 1.**
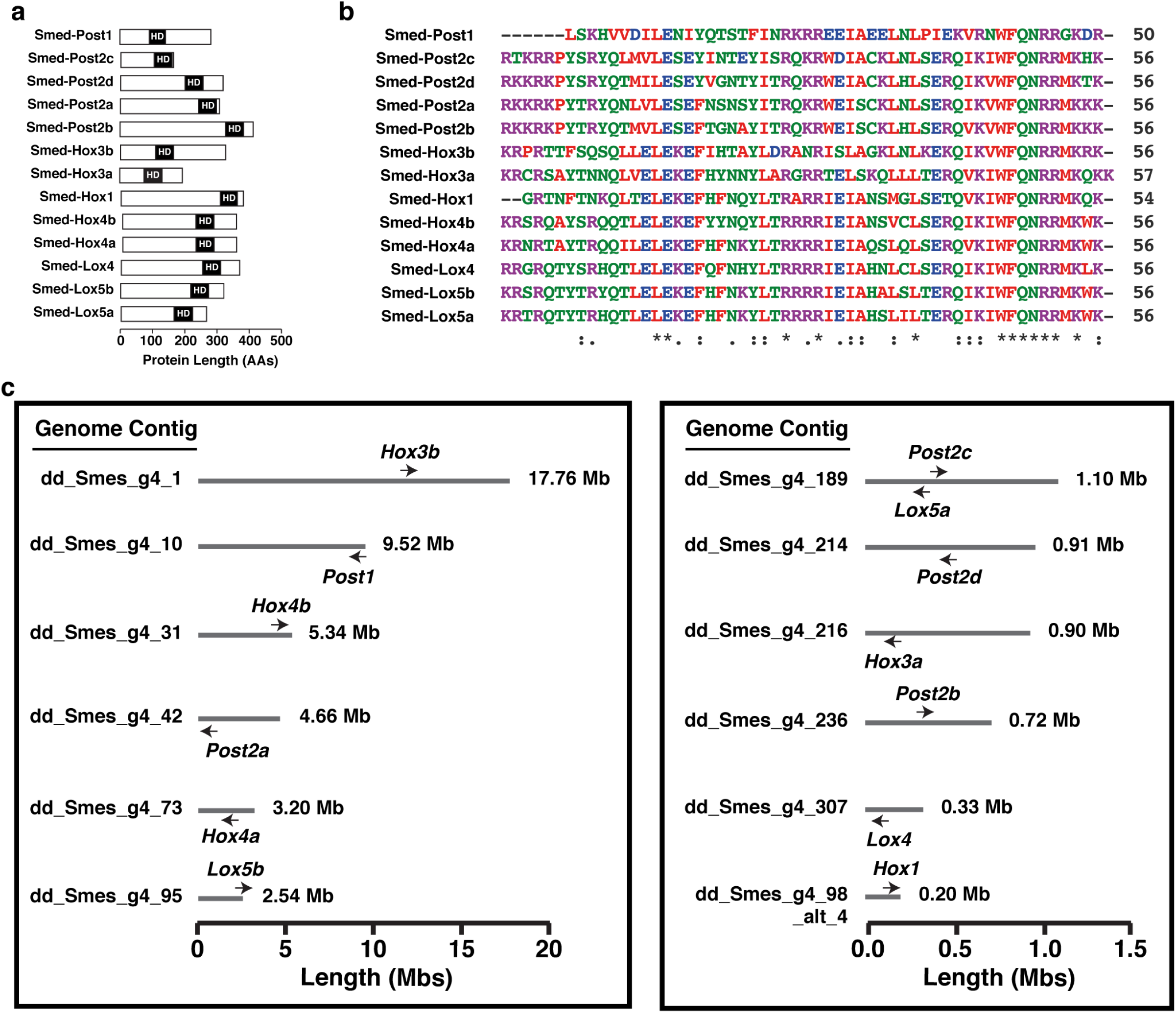
*Hox* genes of the planaria *Schmidtea mediterranea.* **a,** Relative size, hexapeptide motif (HX) location, and conserved homeodomain location (HD) of the 13 planarian Hox family proteins of *Schmidtea mediterranea*. **b,** Sequence alignment of the homeodomains of the 13 planarian Hox family proteins. **c**, Diagram of the genomic distribution, location, and directionality of the 13 planarian *Hox* family genes.

**Extended Data Figure 2.**
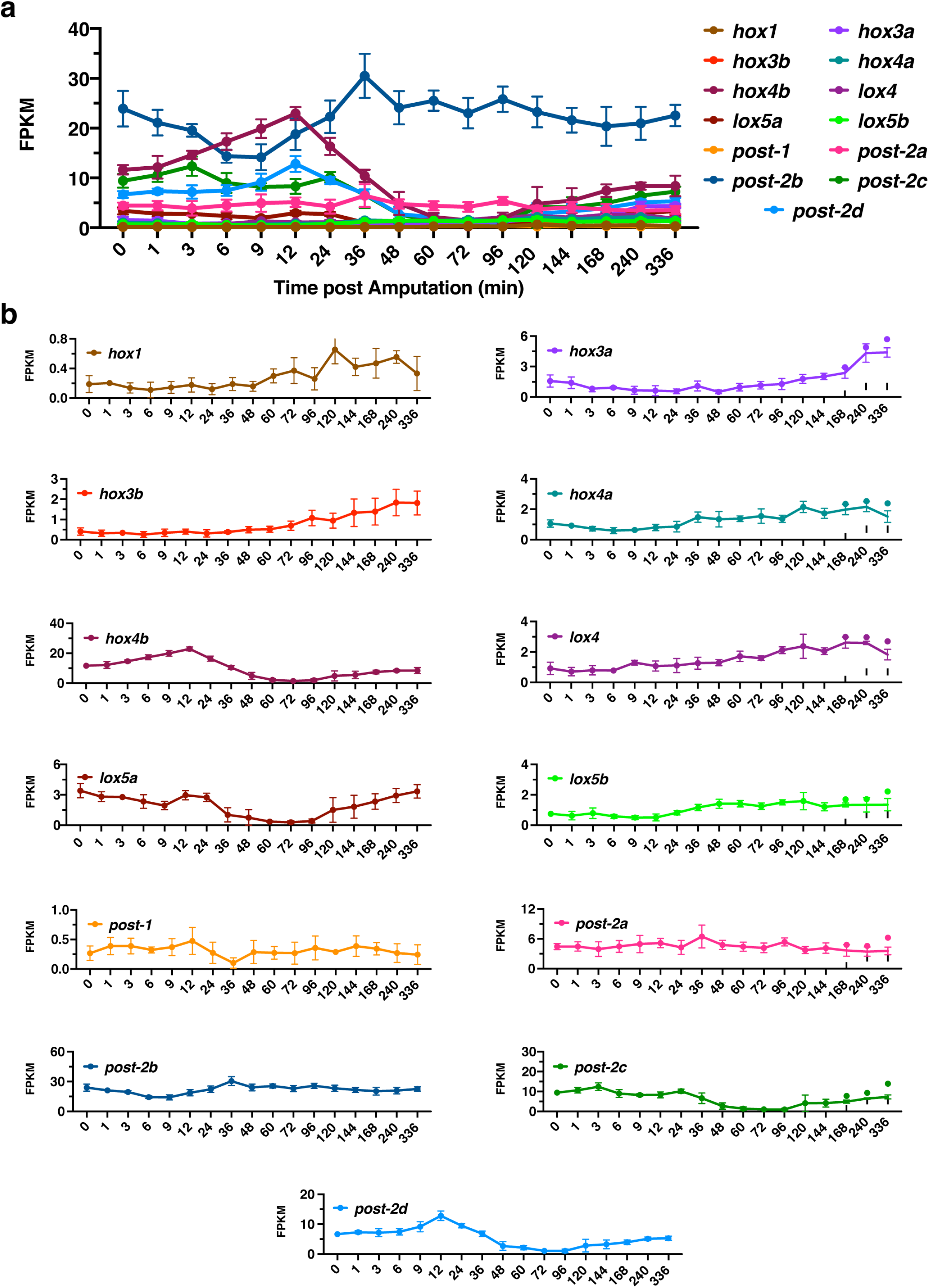
*Hox* gene expression dynamics during planarian regeneration. **a-b,** Time courses of the mRNA expression of (**a**) all 13 *Hox* genes or (**b**) each individual *Hox* gene during the regeneration of laterally amputated tissue fragments. Expression levels plotted as FPKM from RNAseq analysis.

**Extended Data Figure 3.**
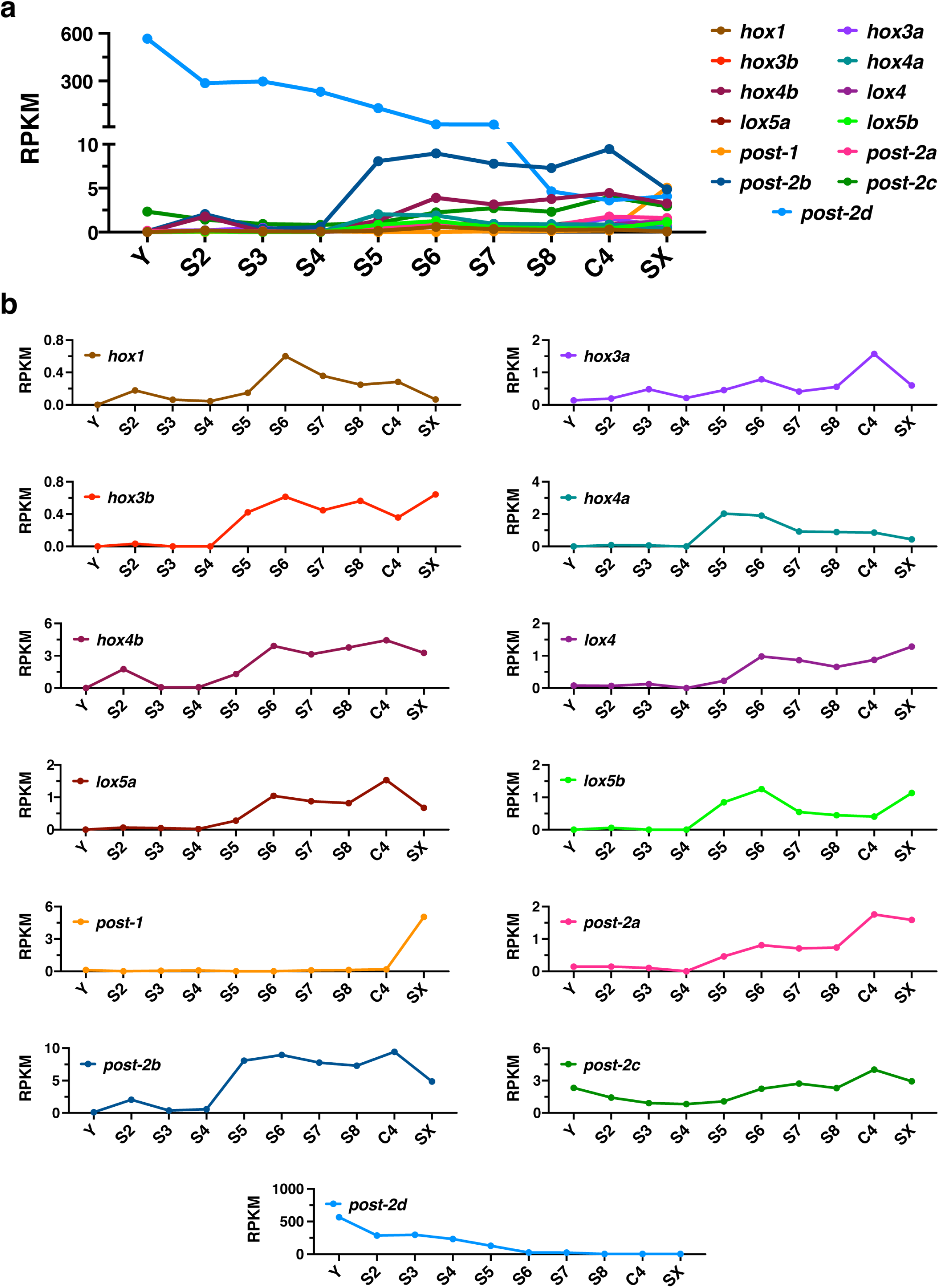
*Hox* gene expression dynamics from planarian embryogenesis to adulthood. **a-b,** Time courses of the mRNA expression of (**a**) all 13 *Hox* genes or (**b**) each individual *Hox* gene at each stage of embryogenesis of the sexual strain and at the adult stage of the sexual and asexual strains. Expression levels plotted as RPKM from RNAseq analysis (Y= egg yolk, S2-S8 = embryonic stages 2–8, C4= adult asexual planaria, SX= adult sexual planaria).

**Extended Data Figure 4.**
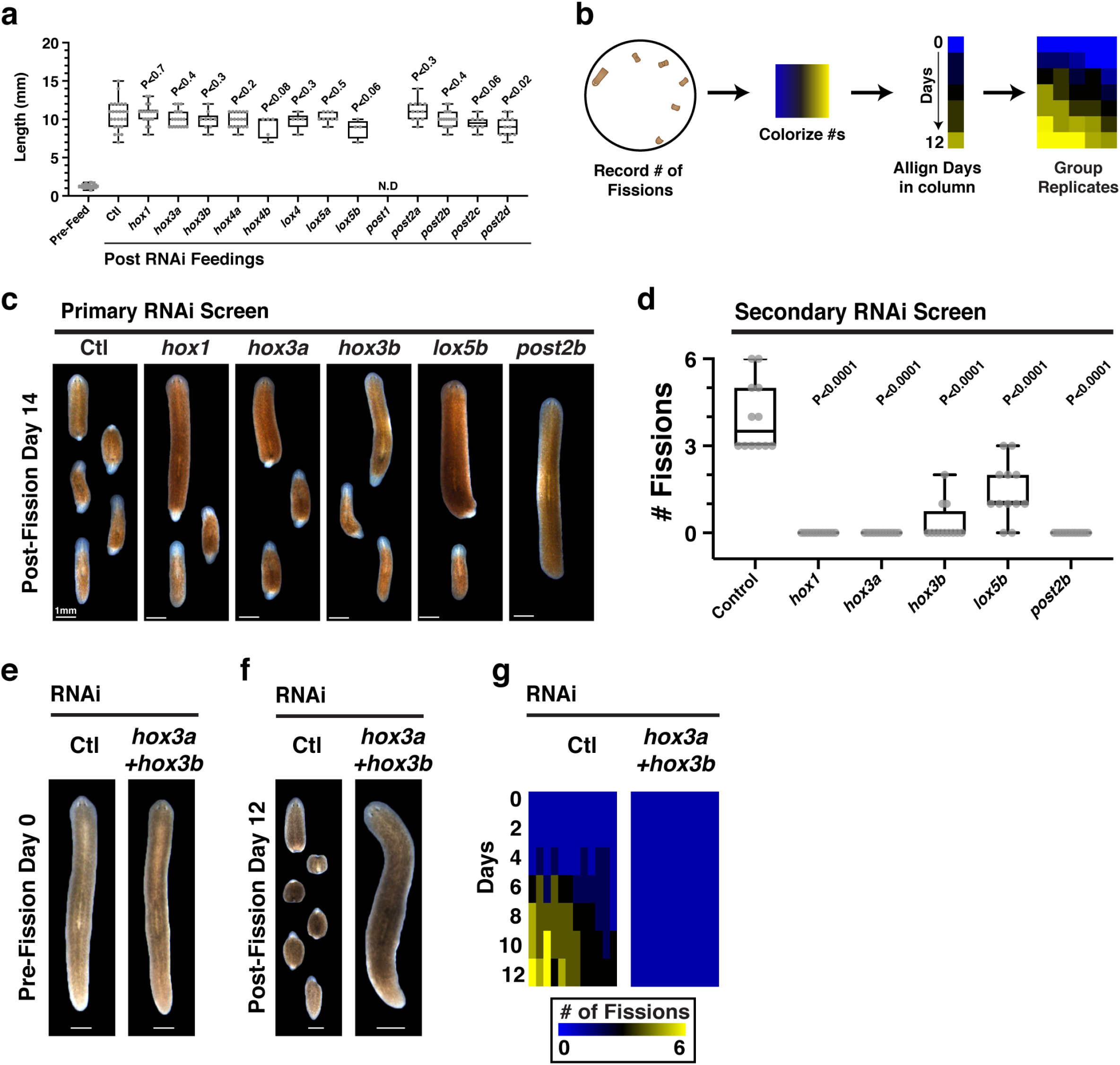
A subset of planarian *Hox* genes regulate fission. **a,** Size distribution of planaria prior to and following 9 purified dsRNA feedings targeting *Hox* genes. **b,** Diagram of data visualization. The number of cumulative fissions for each day was converted to a heat color code. Daily fissions for each individual worm were aligned in ascending order along the y-axis. The average score of each column is calculated and used to sort individual worms in descending order along the x-axis. The result is a heat-map visualization of fission activity across replicates. **c,** Representative images of animals from primary *Hox* RNAi screen on day 14 of the fission assay (n= 18 animals). **d,** Plot depicts fission progeny number on day 12 of secondary RNAi screen (n=12 animals). **e-f,** Representative images of animals of control and *hox3a*+*hox3b* RNAi animals (**e**) prior to and (**f**) 12 days after fission induction (n=12, 17 bacterial RNAi feedings). **g,** Heatmaps depicting cumulative fissions over time for individual worms following control and *hox3a* + *hox3b* RNAi-treatment (n=12 animals). p-value calculated by Welch’s two-tailed t-test versus corresponding control. Scale, 1mm. Note: Control RNAi animals from **e-g** are from the same experiment as **Fig 1 e-f**.

**Extended Data Figure 5.**
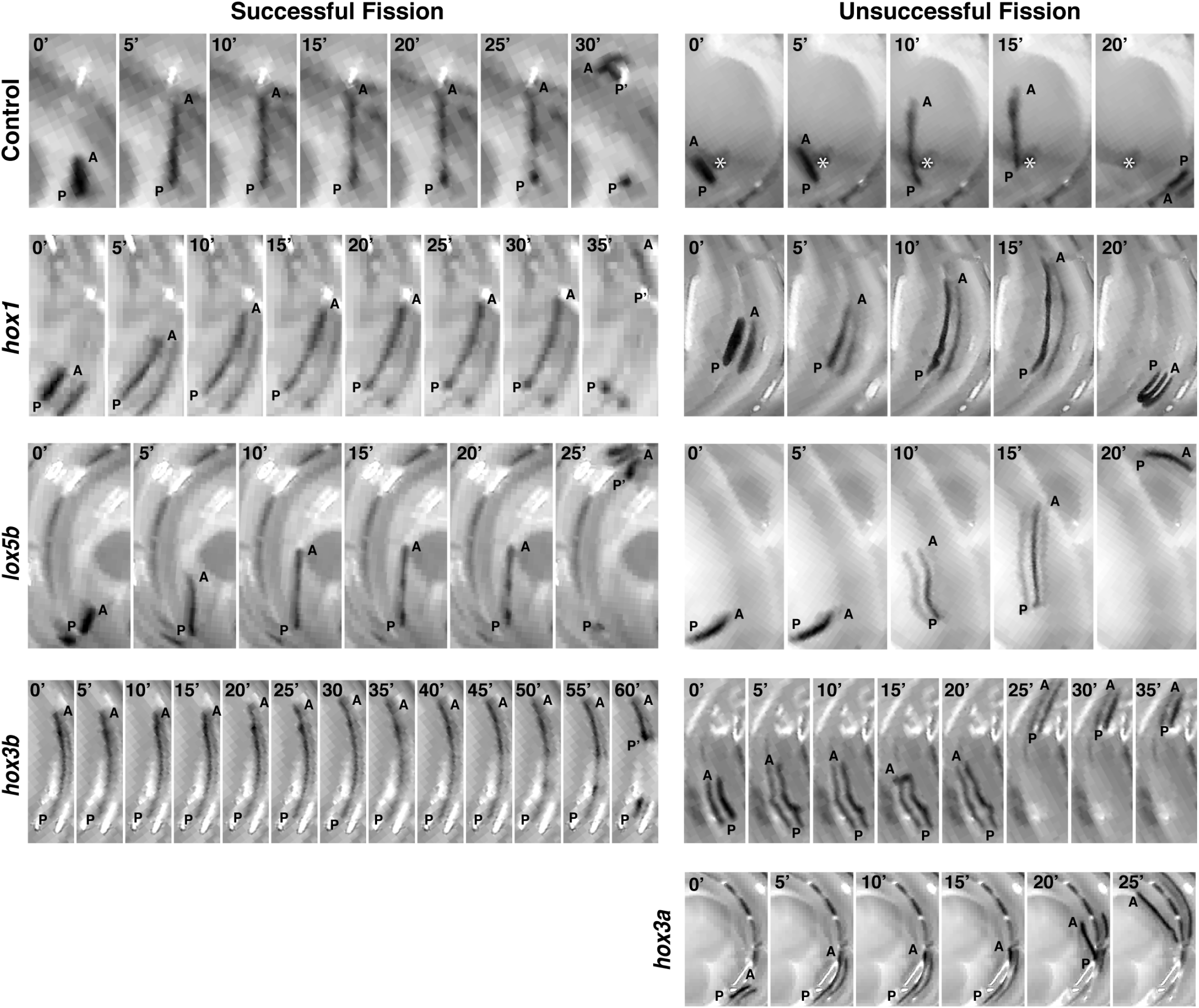
Time-lapse Images of Successful and Unsuccessful fissions following *Hox* gene RNAi-treatment. Representative time courses of fission attempts in RNAi-treated animals (A= anterior, P=pre-fission posterior, and P’=post-fission posterior). Animals fed purified dsRNA food 8 times.

**Extended Data Figure 6.**
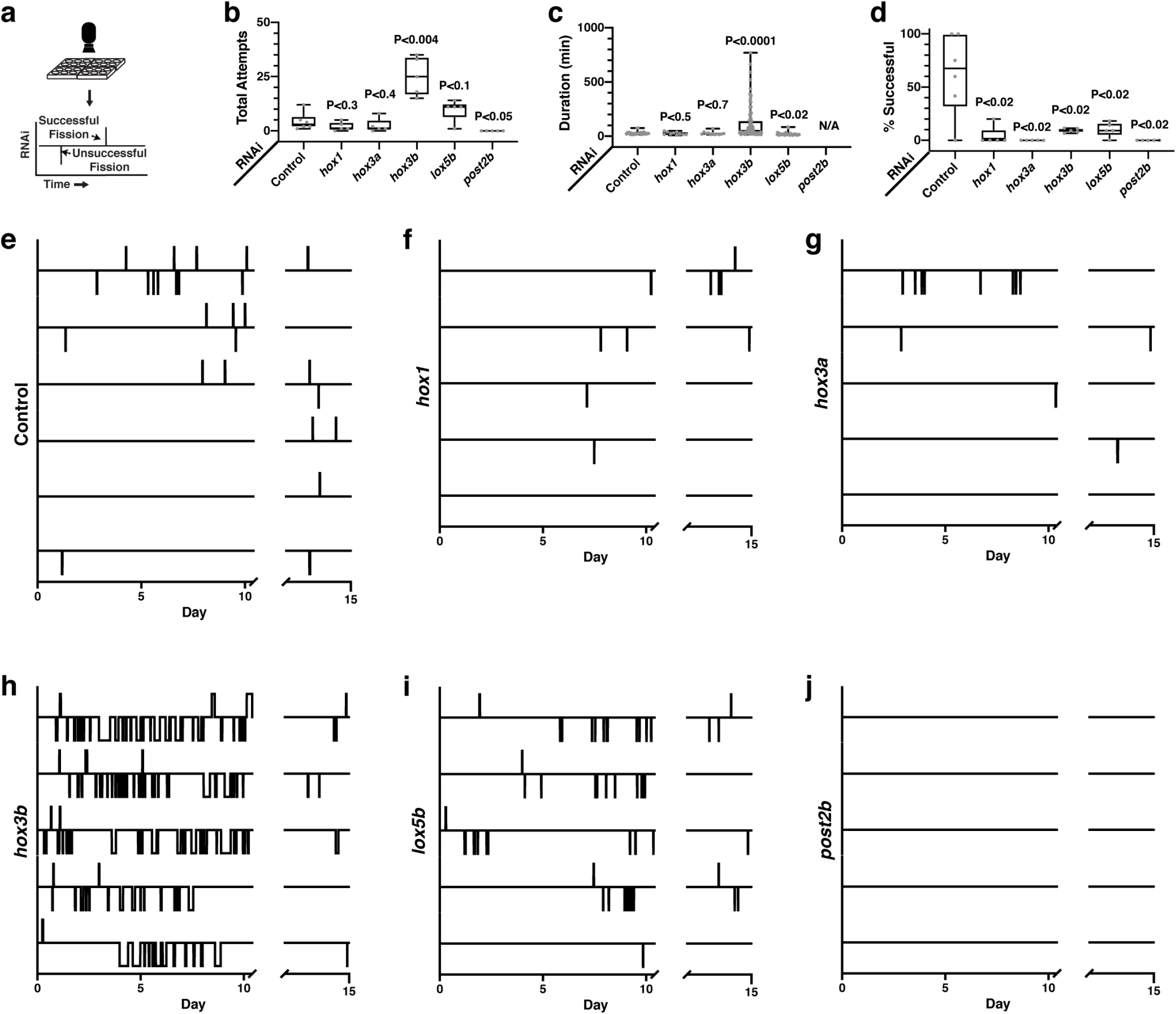
*Hox* genes regulate the frequency, duration, and success of fission behavior. **a,** Schematic of webcam live imaging data visualization. Timeline depicts successful (upward displacement) and unsuccessful fissions (downward displacements). **b-d,** Plots of the (**b**) total, (**c**) duration of, and (**d**) percentage of successful fission attempts following *Hox* RNAi-treatment (n=5-6 animals, p-value calculated by Welch’s two-tailed t-test). **e-j,**fission activity timelines of (**e**) control, (**f**) *hox1*, (**g**) *hox3a*, (**h**) *hox3b*, (**i**) *lox5b*, and (**j**) *post2b* RNAi-treated animals. Animals fed purified dsRNA food 8 times. Discontinuity in timeline due to gap in image acquisition from technical error.

**Extended Data Figure 7.**
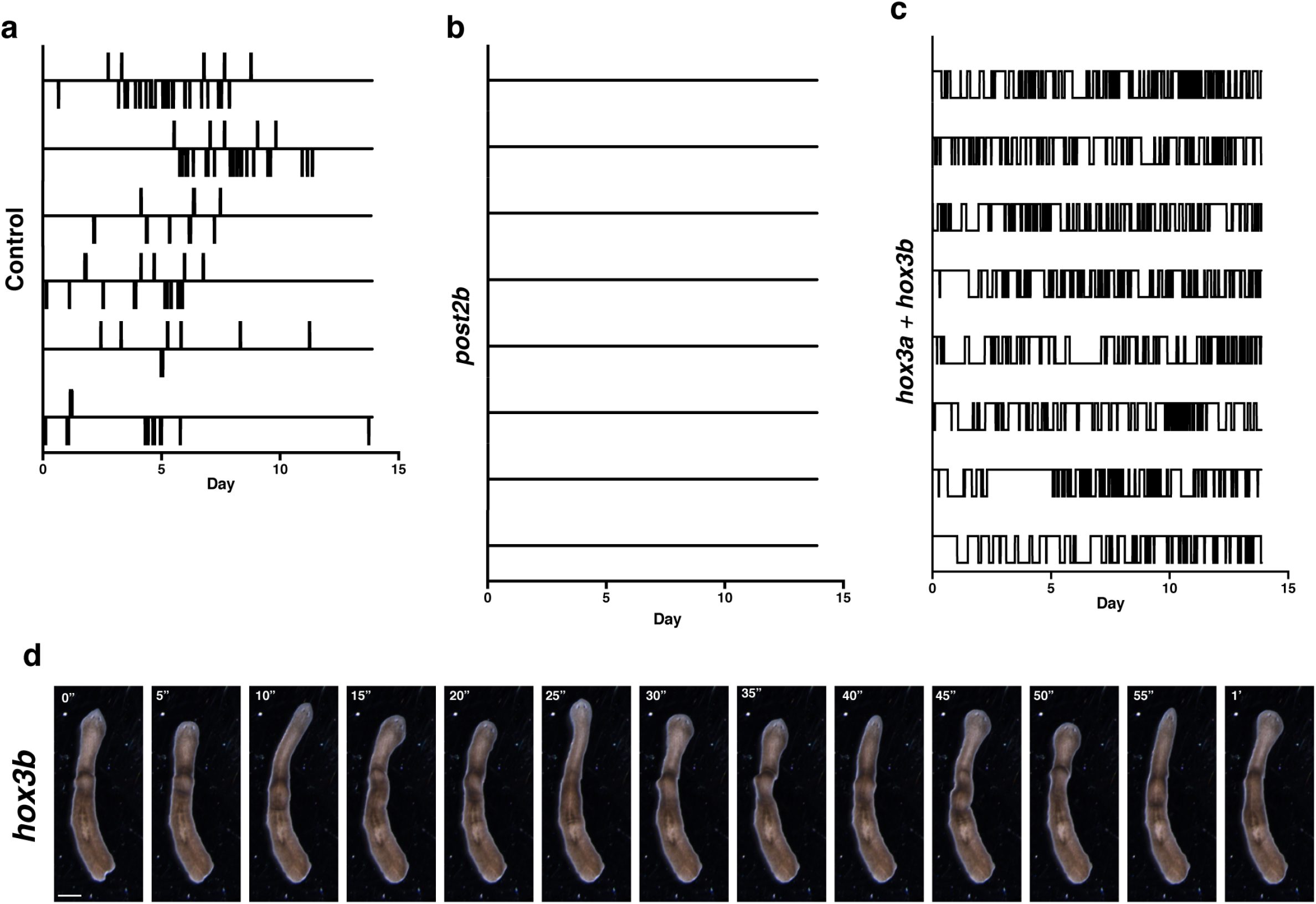
*Post2b* and *hox3* mediate opposing regulation of fission behavior. **a-c,** Fission activity timelines of (**a**) control, (**b**) *post2b*, (**c**) *hox3a* + *hox3b* RNAi-treated animals (n=6-8 animals). **d,**Representative images of *hox3b* RNAi-treated animals stalled in Phase 1 (n=3/10 animals). Animals fed bacterial RNAi food 17 times. Scale, 1mm.

**Extended Data Figure 8.**
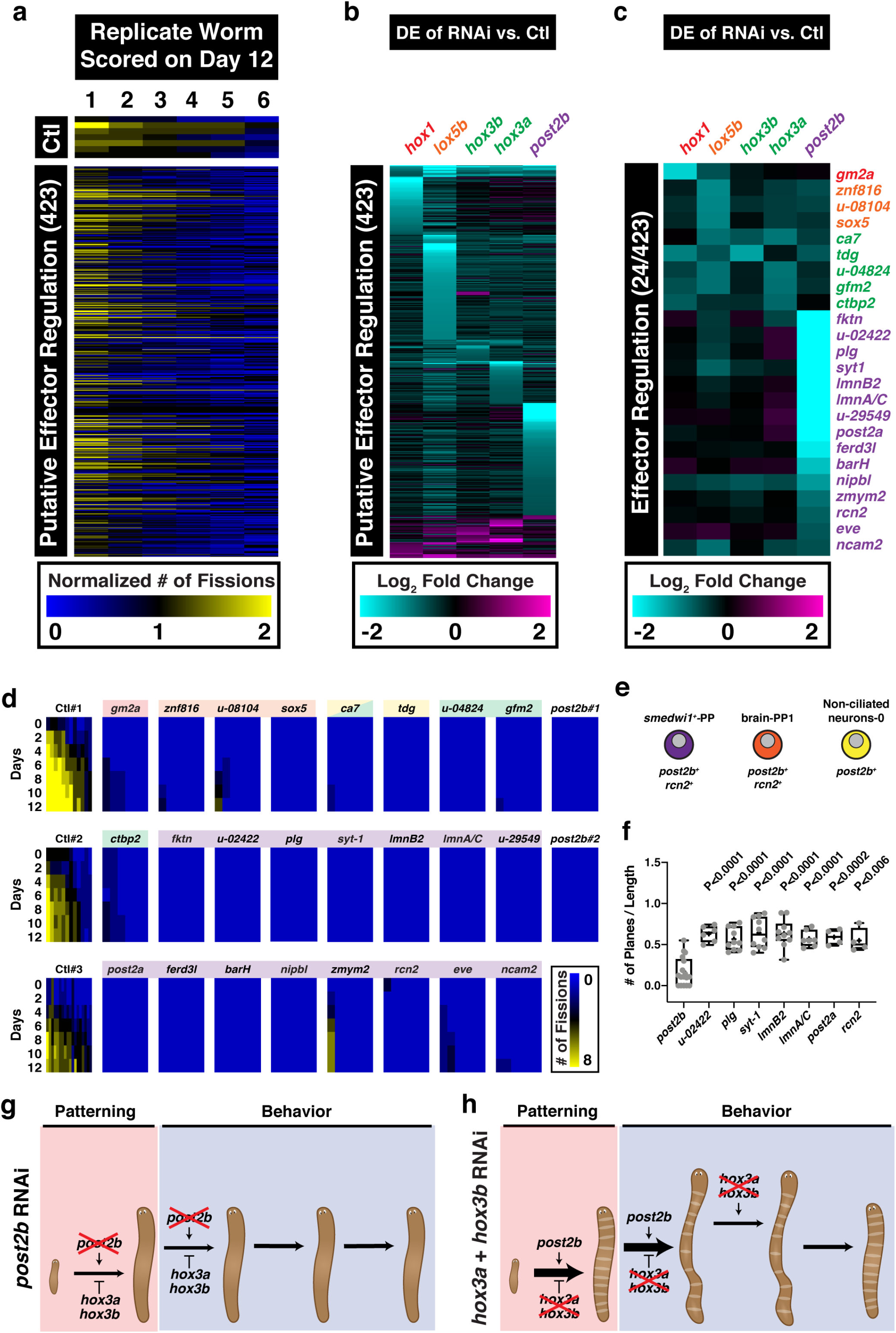
A screen for downstream effectors of *Hox* genes. **a,** Heatmap depicting resultant fission activity following RNAi individually targeted against 423 putative effector genes from the DEGs of the *Hox* RNAi RNAseq (fed 16–19 times with bacterial RNAi food). Heatmap depicts normalized fission number on Day 12 across six RNAi-treated worms. **b,** Heatmap of the corresponding DE of each of the 423 putative fission effectors following RNAi of Hox genes. **c,** Heatmap of the corresponding DE of each of the 24 fission effectors following RNAi of Hox genes. **d**, Heatmaps of the number of fissions over time following RNAi of fission effectors (n=6-18 animals). **e,** Cell population specific enrichment *post2b* and *rcn2* in the *smedwi*^*+*^, brain, and non-cilliated neuronal cell lineages (summary of data from the cell type transcriptome atlas Fincher et al, digiworm.mit.edu). See Figure 4 for epidermal and parenchymal cell lineages (10/11 *post2b* effectors). **f,** Plot of the number of fission planes/animal length following *post2b* effector RNAi treatment (n=4-17 animals, pool of 1–2 independent experiments). p-values calculated by Welch’s two-tailed t-test. **g-h,** Diagram summarizing the patterning and behavioral phenotypes following (**g**) *post2b* and (**h**) *hox3a*+*hox3b* RNAi.

## Tables

**Table 1. Downstream effectors mediating Hox gene regulation of asexual reproduction as identified by RNAseq analysis and RNAi screening.**

## Videos

**Video S1. Fission behavior in Control RNAi animals (Time-lapse A).**

Representative movie of fission behavior in control (*unc-22*) RNAi animals. Worms were placed in 6-well dishes with cameras mounted above the plates and images acquired every 5 minutes for 15 days.

**Video S2. Fission behavior in *hox1* RNAi animals (Time-lapse A).**

Representative movie of fission behavior in *hox1* RNAi animals. Worms were placed in 6-well dishes with cameras mounted above the plates and images acquired every 5 minutes for 15 days.

**Video S3. Fission behavior in *hox3a* RNAi animals (Time-lapse A).**

Representative movie of fission behavior in *hox3a* RNAi animals. Worms were placed in 6-well dishes with cameras mounted above the plates and images acquired every 5 minutes for 15 days.

**Video S4. Fission behavior in *hox3b* RNAi animals (Webcam Time-lapse A).**

Representative movie of fission behavior in *hox3b* RNAi animals. Worms were placed in 6-well dishes with cameras mounted above the plates and images acquired every 5 minutes for 15 days.

**Video S5. Fission behavior in *lox5b* RNAi animals (Webcam Time-lapse A).**

Representative movie of fission behavior in *lox5b* RNAi animals. Worms were placed in 6-well dishes with cameras mounted above the plates and images acquired every 5 minutes for 15 days.

**Video S6. Fission behavior in Control RNAi animals (Webcam Time-lapse B).**

Representative movie of fission behavior in control (*unc-22*) RNAi animals. Worms were placed in 6-well dishes with cameras below the plates and images acquired every 5 minutes for 15 days.

**Video S7. Fission behavior in *post2b* RNAi animals (Webcam Time-lapse B).**

Representative movie of fission behavior in *post2b* RNAi animals. Worms were placed in 6-well dishes with cameras below the plates and images acquired every 5 minutes for 15 days.

**Video S8. Fission behavior in *hox3a+hox3b* RNAi animals (Webcam Time-lapse B).**

Representative movie of fission behavior in *hox3a+hox3b* RNAi animals. Worms were placed in 6-well dishes with cameras below the plates and images acquired every 5 minutes for 15 days.

**Video S9. Phase I peristalsis in***hox3a+hox3b* **RNAi animals.**

Representative movie of peristalsis after fission initiation in *hox3a+hox3b* RNAi animals. Images acquired at 10 frames per second and played at 5X speed.

**Video S10. Phase I peristalsis in *hox3b* RNAi animals.**

Representative movie of peristalsis after fission initiation in *hox3b* RNAi animals. Images acquired at 10 frames per second and played at 5X speed.

## References

1 Gaunt, S. J. Hox cluster genes and collinearities throughout the tree of animal life. Int J Dev Biol 62, 673–683, doi:10.1387/ijdb.180162sg (2018).

2 Rux, D. R. & Wellik, D. M. Hox genes in the adult skeleton: Novel functions beyond embryonic development. Dev Dyn 246, 310–317, doi:10.1002/dvdy.24482 (2017).

3 Bradaschia-Correa, V. et al. Hox gene expression determines cell fate of adult periosteal stem/progenitor cells. Sci Rep 9, 5043, doi:10.1038/s41598-019-41639-7 (2019).

4 Arnold, C. P., Benham-Pyle B. W., Lange, J. J., Wood, C. J., Sánchez Alvarado, A. Wnt and TGFβ coordinate growth and patterning to regulate size-dependent behavior. Nature (2019).

5 Malinowski, P. T. et al. Mechanics dictate where and how freshwater planarians fission. Proc Natl Acad Sci U S A 114, 10888–10893, doi:10.1073/pnas.1700762114 (2017).

6 He, S. et al. An axial Hox code controls tissue segmentation and body patterning in Nematostella vectensis. Science 361, 1377–1380, doi:10.1126/science.aar8384 (2018).

7 Currie, K. W. et al. HOX gene complement and expression in the planarian Schmidtea mediterranea. Evodevo 7, 7, doi:10.1186/s13227-016-0044-8 (2016).

8 Zeng, A. et al. Prospectively Isolated Tetraspanin(+) Neoblasts Are Adult Pluripotent Stem Cells Underlying Planaria Regeneration. Cell 173, 1593–1608 e1520, doi:10.1016/j.cell.2018.05.006 (2018).

9 Davies, E. L. et al. Embryonic origin of adult stem cells required for tissue homeostasis and regeneration. Elife 6, doi:10.7554/eLife.21052 (2017).

10 Tewari, A. G., Owen, J. H., Petersen, C. P., Wagner, D. E. & Reddien, P. W. A small set of conserved genes, including sp5 and Hox, are activated by Wnt signaling in the posterior of planarians and acoels. PLoS Genet 15, e1008401, doi:10.1371/journal.pgen.1008401 (2019).

11 Wurtzel, O., Oderberg, I. M. & Reddien, P. W. Planarian Epidermal Stem Cells Respond to Positional Cues to Promote Cell-Type Diversity. Dev Cell 40, 491–504 e495, doi:10.1016/j.devcel.2017.02.008 (2017).

12 Svingen, T. & Tonissen, K. F. Hox transcription factors and their elusive mammalian gene targets. Heredity (Edinb) 97, 88–96, doi:10.1038/sj.hdy.6800847 (2006).

13 Jones, F. S., Prediger, E. A., Bittner, D. A., De Robertis, E. M. & Edelman, G. M. Cell adhesion molecules as targets for Hox genes: neural cell adhesion molecule promoter activity is modulated by cotransfection with Hox-2.5 and −2.4. Proc Natl Acad Sci U S A 89, 2086–2090, doi:10.1073/pnas.89.6.2086 (1992).

14 Fincher, C. T., Wurtzel, O., de Hoog, T., Kravarik, K. M. & Reddien, P. W. Cell type transcriptome atlas for the planarian Schmidtea mediterranea. Science 360, doi:10.1126/science.aaq1736 (2018).

15 Philippidou, P. & Dasen, J. S. Hox genes: choreographers in neural development, architects of circuit organization. Neuron 80, 12–34, doi:10.1016/j.neuron.2013.09.020 (2013).

16 Honore, B. The rapidly expanding CREC protein family: members, localization, function, and role in disease. Bioessays 31, 262–277, doi:10.1002/bies.200800186 (2009).

17 Bello, O. D. et al. Synaptotagmin oligomerization is essential for calcium control of regulated exocytosis. Proc Natl Acad Sci U S A 115, E7624–E7631, doi:10.1073/pnas.1808792115 (2018).

18 Verzi, M. P. et al. N-twist, an evolutionarily conserved bHLH protein expressed in the developing CNS, functions as a transcriptional inhibitor. Dev Biol 249, 174–190, doi:10.1006/dbio.2002.0753 (2002).

19 Reig, G., Cabrejos, M. E. & Concha, M. L. Functions of BarH transcription factors during embryonic development. Dev Biol 302, 367–375, doi:10.1016/j.ydbio.2006.10.008 (2007).

20 Doe, C. Q., Smouse, D. & Goodman, C. S. Control of neuronal fate by the Drosophila segmentation gene even-skipped. Nature 333, 376–378, doi:10.1038/333376a0 (1988).

21 Coffinier, C., Fong, L. G. & Young, S. G. LINCing lamin B2 to neuronal migration: growing evidence for cell-specific roles of B-type lamins. Nucleus 1, 407–411, doi:10.4161/nucl.1.5.12830 (2010).

22 Oyston, L. J. et al. Neuronal Lamin regulates motor circuit integrity and controls motor function and lifespan. Cell Stress 2, 225–232, doi:10.15698/cst2018.09.152 (2018).

23 Alexander, T., Nolte, C. & Krumlauf, R. Hox genes and segmentation of the hindbrain and axial skeleton. Annu Rev Cell Dev Biol 25, 431–456, doi:10.1146/annurev.cellbio.042308.113423 (2009).

24 Reilly, M. B., Cros, C., Varol, E., Yemini, E. & Hobert, O. Unique homeobox codes delineate all the neuron classes of C. elegans. Nature 584, 595–601, doi:10.1038/s41586-020-2618-9 (2020).

25 Petersen, C. P. & Reddien, P. W. Wnt signaling and the polarity of the primary body axis. Cell 139, 1056–1068, doi:10.1016/j.cell.2009.11.035 (2009).

26 Sköld HN, O. M., Sköld M, Åkesson B. Stem cells in asexual reproduction of marine invertebrates. Vol. Stem cells in marine organisms 105–37 (Springer Netherlands, 2009).

27 Arnold, C. P. et al. Pathogenic shifts in endogenous microbiota impede tissue regeneration via distinct activation of TAK1/MKK/p38. Elife 5, doi:10.7554/eLife.16793 (2016).

28 Dobin, A. et al. STAR: ultrafast universal RNA-seq aligner. Bioinformatics 29, 15–21, doi:10.1093/bioinformatics/bts635 (2013).

29 Robinson, M. D., McCarthy, D. J. & Smyth, G. K. edgeR: a Bioconductor package for differential expression analysis of digital gene expression data. Bioinformatics 26, 139–140, doi:10.1093/bioinformatics/btp616 (2010).

30 Benjamini, Y. & Hochberg, Y. Controlling the False Discovery Rate: A Practical and Powerful Approach to Multiple Testing. Journal of the Royal Statistical Society: Series B (Methodological) 57, 289–300, doi:10.1111/j.2517-6161.1995.tb02031.x (1995).

31 El-Gebali, S. et al. The Pfam protein families database in 2019. Nucleic Acids Res 47, D427–D432, doi:10.1093/nar/gky995 (2019).

